# Assessing the viability of biochemical networks across planets

**DOI:** 10.1101/607531

**Authors:** Harrison B. Smith, Alexa Drew, Sara I. Walker

## Abstract

The concept of the origin of life implies that initially, life emerged from a non-living medium. If this medium was Earth’s geochemistry, then that would make life, by definition, a geochemical process. The extent to which life on Earth today could subsist outside of the geochemistry from which it is embedded is poorly quantified. By leveraging large biochemical datasets in conjunction with planetary observations and computational tools, this research provides a methodological foundation for the quantitative assessment of our biology’s viability in the context of other geospheres. Investigating a case study of alkaline prokaryotes in the context of Enceladus, we find that the chemical compounds observed on Enceladus thus far would be insufficient to allow even these extremophiles to produce the compounds necessary to sustain a viable metabolism. The environmental precursors required by these organisms provides a map for the compounds which should be prioritized for detection in future planetary exploration missions. The results of this framework have further consequences in the context of planetary protection, and hint that forward contamination may prove infeasible without meticulous intent.

## Introduction

It is probable that the geochemical process known as life had already commenced when today’s oldest minerals began to crystallize. While there is widely accepted evidence that the process of life has been present on Earth continuously for the past 3.4Gy [1], the lack of evidence prior to this date has more to do with the paucity of fossil-preserving rocks than concrete evidence of life’s absence [14, 32]. Despite the biosphere’s apparent interminable coexistence with the geosphere, there remain many open questions on the matter of life persisting in Earth’s absence [3, 35], not to mention the questions of Earth persisting in life’s absence [21, 23, 24]. For example, Visionaries dream of terraforming planets while program officers fret over “contaminating” them [25, 31, 34]. While the terraformers tend to believe that seeding another planet would require careful human or robotic (and usually Earth-assisted) cultivation, planetary protection officers take the more conservative stance that a small, semi-sterilized spacecraft of Earth origin could cause life to spill onto a planet in the same way that a small perturbation to a super cooled liquid would cause the entire volume to quickly crystallize. In both cases, there is the predominately implicit assumption that Earth-life would be viable outside of the Earth.

When life is viewed as a geologic process, this is a somewhat surprising assumption. In the words of Morowitz et al., “the metabolic character of life is a planetary phenomenon, no less than the atmosphere, hydrosphere, or geosphere” [30]. If this “metabolic character of life” is truly a planetary phenomenon, does that imply that life is inextricable from the planet through which it emerged? Or is it possible that an infinitesimal component of our biosphere—a sliver of a sliver of Earth’s biochemical diversity captured in a few species—could be enough to imbue another world with Earth’s vitality?

To begin to address these questions, we must first lay the framework for determining the environmental conditions required for a species to produce or acquire the chemical compounds necessary to yield a viable metabolism. For this, we utilize the network expansion method [13]: an organism can catalyze a reaction only if it has access to the necessary substrates. The initial substrates, called the *seed set*, are the compounds available to the organism from the environment. Initially, these are the only compounds in the organism’s *network*—an abstract representation of the biochemistry able to be utilized by the organism with the given compounds. The organism catalyzes all the reactions it can based on the compounds available in its network, and then adds the new compounds it can generate to its network. This process proceeds iteratively until the organism can produce no new compounds. The state of the organism’s network when expansion ceases is referred to as the organism’s *scope*—and it contains all of the compounds which can be synthesized by an organism, plus the compounds provided by the environment (the seed set).

While there are other methods which can be used to computationally assess organismal viability, relying on some combination of integer linear programming, kinetic modeling using differential equations, elementary mode analysis, and flux balance analysis (FBA), they require catalytic rates which are difficult to acquire and sparsely catalogued, or a curated list of stoichiometrically balanced reactions [27]. FBA is perhaps the most common method for assessing organismal viability, and operates by solving for the relative fluxes of reactions needed in order for steady state production of compounds identified necessary for organismal growth. Despite FBA requiring more constrained information and computational resources, network expansion has been shown to give near identical results for identifying compounds produced (the network scope) [22, 27].

Network expansion models have been used to explore the scope of chemicals accessible to biology across space and time on Earth, and how changing environments and changing biochemical networks impact one another [2]. For example, the models have been utilized to identify how oxygen drastically altered life’s biochemical networks during the great oxygenation event [33]; how biochemistry differed before phosphorous was widely available [10]; how organismal scopes vary across the tree of life [2, 7]; and how organismal metabolic variability is impacted both in the presence of diverse environments and the presence of other species [8].

We propose using network expansions to address the question of life’s viability amongst other planetary chemistries in two fundamental ways: For a set of organisms and a set of planetary environments, how many target substrates can each organism produce across the environments? The inverse question—For a set of organisms and a set of planetary environments, what chemical seed sets must be provided in order to produce the substrates which are necessary to the organism’s viability?

We work through a case study of this framework to determine the viability of varying Earth organisms within Enceladus’s planetary context. Because Enceladus has an ocean with high pH (11-12) [9], we choose to focus on the viability of prokaryotic alkaliphiles. Because other environmental factors are less well constrained, and parameters like temperature and salinity could vary substantially across locations, we do not place any further restrictions on the organismal metabolisms that we run network expansions on [28]. We show that based on the compounds we currently know to be present in Enceladus’s subsurface ocean [37], none of the analyzed organismal metabolisms are viable. In order to verify that this is not solely due to the lack of phosphate, a prominent bioessential compound on Earth which has not been detected on Enceladus (likely due to Cassini instrument detection thresholds), we show that adding phosphate as a seed compound still results in no viable organisms. Using an algorithm developed to solve the inverse network expansion problem [12], we identify minimal sets of substrates that satisfy the requirements of what these alkaliphilic organisms would have to acquire externally in order to produce the target substrates. We find that these organisms tend to require complex molecules and coenzymes, lowering the likelihood that the organisms could be viable on Enceladus, given their lack of detection. Nonetheless, when the full catalytic repertoire of Earth’s biosphere is available, we find that nearly all target substrates are able to be synthesized from a seed set consisting only of the compounds currently observed on Enceladus (plus phosphate). Although these reactions are not the product of organisms which are solely alkaliphilic, these results hint that forward contamination from individuals may be much less concerning than contamination by a microbial ecosystem which can emulate the robustness and catalytic capabilities of the biosphere—reinforcing the perspective that the emergence of life on a planet is an extension of the planet’s geosphere [29, 35]. More importantly, by leveraging large biochemical datasets in conjunction with planetary observations and computational tools, this research provides a methodological foundation for the quantitative assessment of our biology’s viability in the context of other geospheres.

## Results

Based on target metabolites necessary for many living organisms, we first sought to determine if the compounds which have thus far been identified on Enceladus were sufficient to produce the target metabolites in a set of organisms which would be viable in an environment with the alkalinity present on Enceladus [9].

We ran the network expansion algorithm on the subset of archaea and bacteria with documented environmental pH in the ranges of 9-11 [16–18], using a seed set of compounds which have been identified on Enceladus from observations aboard Cassini’s Ion and Neutral Mass Spectrometer (INMS) [37] (Table 1).

**Table 1.** Compounds used for Enceladus seed set. All compounds from Waite et al., 2009 [37] that were present in the Kyoto Encyclopedia of Genes and Genomes (KEGG) were included. Phosphate was added to the seed set for additional analyses.

| Name | Formula | KEGG Compound ID |
| --- | --- | --- |
| Water | (H<sub>2</sub>O) | C00001 |
| Carbon Dioxide | (CO<sub>2</sub>) | C00011 |
| Carbon Monoxide | (CO) | C00237 |
| Hydrogen | (H<sub>2</sub>) | C00282 |
| Formaldehyde | (H<sub>2</sub>CO) | C00067 |
| Methanol | (CH<sub>3</sub>OH) | C00132 |
| Ethylene oxide | (C<sub>2</sub>H<sub>4</sub>O) | C06548 |
| Ethanol | (C<sub>2</sub>H<sub>6</sub>O) | C00469 |
| Hydrogen sulfide | (H<sub>2</sub>S) | C00283 |
| Ammonia | (NH<sub>3</sub>) | C00014 |
| Nitrogen | (N<sub>2</sub>) | C00697 |
| Hydrogen Cyanide | (HCN) | C01326 |
| Methane | (CH<sub>4</sub>) | C01438 |
| Acetylene | (C<sub>2</sub>H<sub>2</sub>) | C01548 |
| Ethylene | (C<sub>2</sub>H<sub>4</sub>) | C06547 |
| Propene | (C<sub>3</sub>H<sub>6</sub>) | C11505 |
| Propane | (C<sub>3</sub>H<sub>8</sub>) | C20783 |
| Benzene | (C<sub>6</sub>H<sub>6</sub>) | C01407 |
| Phosphate | (H<sub>3</sub>PO<sub>4</sub>) | C00009 |

We deem an organism or network to be fully viable if, given a set of environmental seed compounds, it has the catalytic repertoire to produce all the compounds in its network which intersect with a pre-defined set of target metabolites. For this study, we adopt the list of target metabolites defined by Freilich et al (2009) [8], (Table 2). In that study, the authors found that the organisms which were found to be viable, based on these target metabolites, accurately predicted the ecological compositions of known environments across many habitats and bacterial metabolisms.

**Table 2.** Compounds in the target metabolite set. Target list adopted from Freilich et al (2009) [8]

| Name | KEGG Compound ID | Name | KEGG Compound ID |
| --- | --- | --- | --- |
| ATP | C00002 | NAD+ | C00003 |
| NADH | C00004 | NADPH | C00005 |
| NADP+ | C00006 | ADP | C00008 |
| UDP | C00015 | FAD | C00016 |
| AMP | C00020 | Acetyl-CoA | C00024 |
| L-Glutamate | C00025 | GDP | C00035 |
| Glycine | C00037 | L-Alanine | C00041 |
| UDP-N-acetyl-D-glucosamine | C00043 | GTP | C00044 |
| L-Lysine | C00047 | L-Aspartate | C00049 |
| Adenosine 3',5'-bisphosphate | C00054 | CMP | C00055 |
| L-Arginine | C00062 | CTP | C00063 |
| L-Glutamine | C00064 | L-Serine | C00065 |
| L-Methionine | C00073 | UTP | C00075 |
| L-Tryptophan | C00078 | L-Phenylalanine | C00079 |
| L-Tyrosine | C00082 | L-Cysteine | C00097 |
| UMP | C00105 | CDP | C00112 |
| Glycerol | C00116 | L-Leucine | C00123 |
| dATP | C00131 | L-Histidine | C00135 |
| GMP | C00144 | L-Proline | C00148 |
| L-Asparagine | C00152 | L-Valine | C00183 |
| L-Threonine | C00188 | 10-Formyltetrahydrofolate | C00234 |
| dCMP | C00239 | Hexadecanoic acid | C00249 |
| Riboflavin | C00255 | dGTP | C00286 |
| Phosphatidylethanolamine | C00350 | dAMP | C00360 |
| dGMP | C00362 | dTMP | C00364 |
| Ubiquinone | C00399 | L-Isoleucine | C00407 |
| dCTP | C00458 | dTTP | C00459 |
| 1,2-Diacyl-sn-glycerol | C00641 | Siroheme | C00748 |
| UDP-N-acetylmuramate | C01050 | Hexadecanoyl-[acp] | C05764 |
| Cardiolipin | C05980 | Diglucosyl-diacylglycerol | C06040 |
| Heme O | C15672 | (2E)-Octadecenoyl-[acp] | C16221 |
| Undecaprenyl-diphospho-... | C05890 |  |  |
| N-acetylmuramoyl-... | C05894 |  |  |
| (N-acetylglucosamine)-L | C05899 |  |  |

### Prokaryotic viability on Enceladus

We find that none of these organisms, across bacteria and archaea, can produce any target metabolites with the few identified organic and inorganic compounds on Enceladus. In fact, they are found to produce only a fraction of the compounds possible given their reaction network (Fig. 1). However, this was not surprising given the lack of detection of any phosphorous containing compounds. Because of this, we repeated the expansion with the addition of phosphate. While this increased the scope of the organismal seed sets, again, no target compounds were able to be produced. Although in the latter case, we note that the organismal scopes increased in size (Fig. 1A).

**Figure 1.**
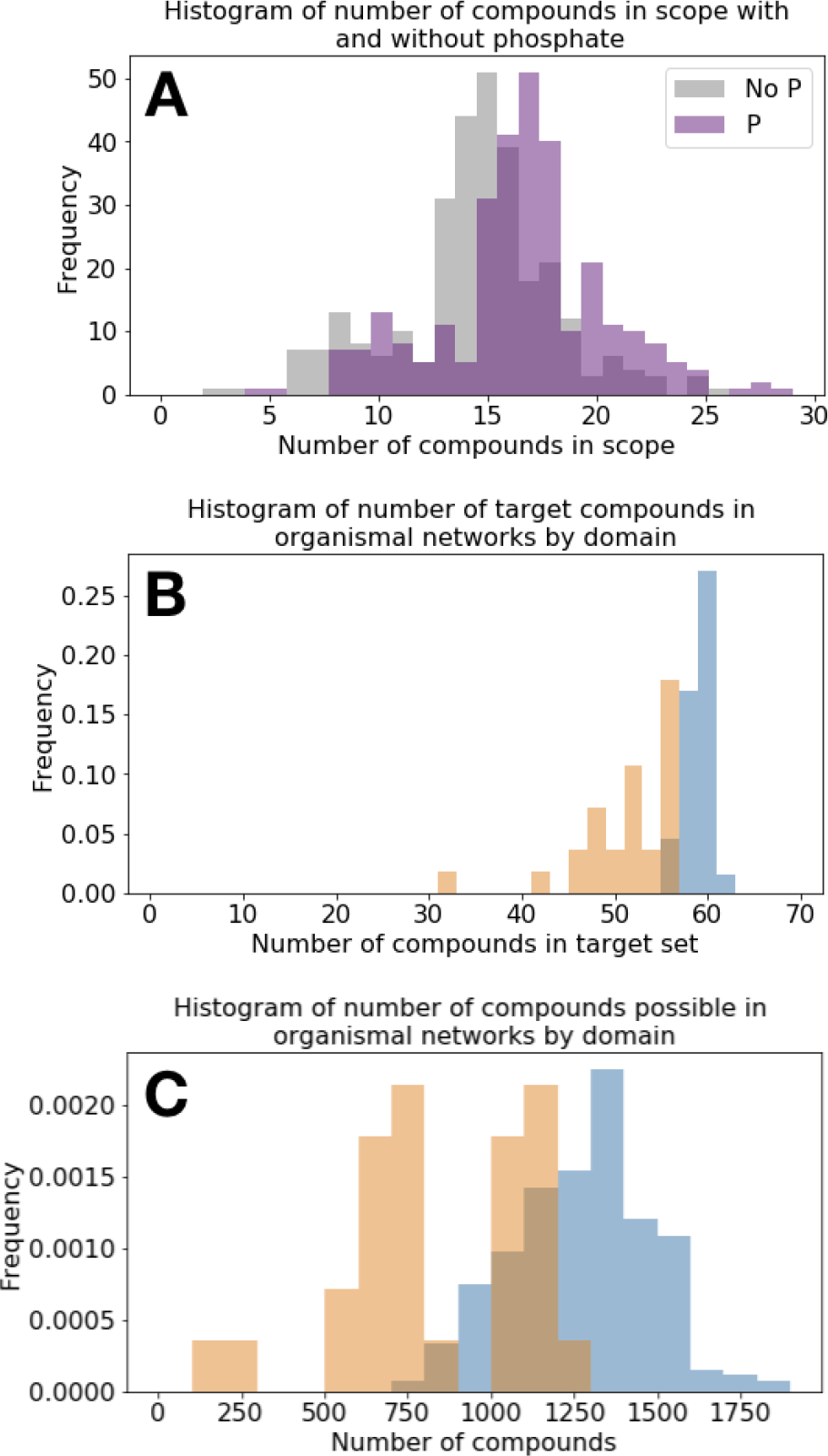
Histograms from the network expansions for prokaryotes using the Enceladus seed set. (A) How the scope size changes for all organisms when adding phosphate to the seed set adopted from Waite et al. [37]. In neither case do any target compounds get produced for any organisms. (B) An overview of the distribution of number of target compounds across all organisms (out of 65 possible based on the target set from Freilich et al. [8]. (C) The maximum theoretical sizes of networks, if scopes were able to take advantage of full organismal reaction networks. Orange bars are for archaea, and blue bars are for bacteria.

### Identifying the compounds necessary to make prokaryotes viable

Running network expansions on pre-established seed sets are useful for determining the set of compounds which can be part of an organism’s scope. However, as we found in the section above, if we are aiming to produce a specific set of target compounds, there is no guarantee that a chosen seed set will do that. For this reason, it is useful to identify an algorithm which can identify the seed set needed to produce a target set, given a reaction network. We thus sought to identify subsets of all compounds involved in each organism’s network which could feasibly produce all the target compounds in that network.

There are three obvious ways to go about this. We could imagine searching for: 1) a single minimal seed set (no subsets of which can produce all target metabolites), 2) the smallest minimal seed set (where there are no sets with fewer elements which can produce all target metabolites), or 3) all minimal seed sets (the set of all sets that can produce all target metabolites).

We chose to identify a subset of all minimal seed sets for the archaea and bacteria under consideration, because finding the smallest minimal seed set is an NP-hard problem (Cottret et al., 2008), and because it would result in only a single environment in which a target set could be produced. Finding any given minimal seed set requires a polynomial-time algorithm, so for computational tractability we chose to identify 100 random minimal seed sets for each of the 28 aforementioned archaea, and for 36 of the aforementioned 266 bacteria. We follow the algorithm described in Handorf et al., 2008 to create random minimal seed sets which attempt to minimize the likelihood of obtaining seed sets with large complex biomolecules where possible (see methods).

We first take an overview of the minimal seed sets we find which produce target compounds for each of the analyzed organisms. We find that the environmental seed sets needed are often smaller in size, but more complex (as quantified by the mean molecular weight of the seed sets needed) (Fig. 2). This is especially true for the bacteria, while for archaea the seed sets tend to be composed both of more complex molecules and more of them. Interestingly, there no seeds identified which require more than four of the compounds which have been identified as part of the Enceladus seed set.

**Figure 2.**
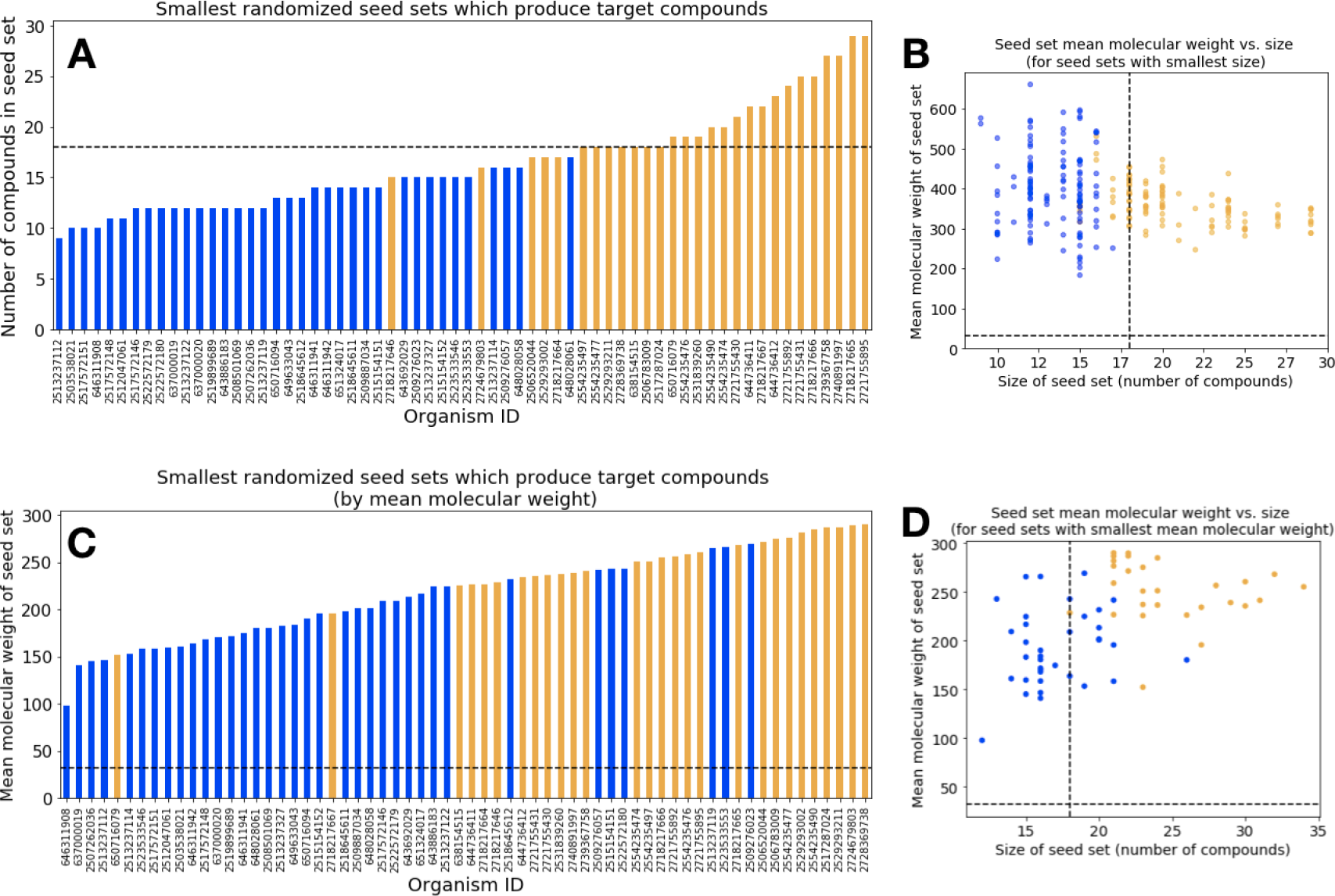
Characteristics of minimal seed sets which produce target metabolites. (A) A rank ordered plot of the smallest minimal seed sets, by number of compounds involved in each seed set. The mean molecular weights of the smallest seed sets, by size, of the seed sets with the smallest size. Note that many organisms have multiple minimal seed sets of the same size, but of different mean molecular weights. (C) A rank ordered plot of the smallest minimal seed sets, by weight. (D) The mean molecular weights of the smallest seed sets, by size, of the seed sets with smallest mean molecular weight. Orange bars are archaea, and blue lines are bacteria, with each organism represented on the x-axis. The black dashed lines in each case shows the size and weight values for the Enceladus seed set.

Next we look at how similar each of the 100 minimal seeds sets for each organism are to one another. We find that across all organisms, the archaea seed sets tend to have more self-similarity compared to the bacteria. Two archaea share about a quarter of the compounds across all their seed sets, on average (Fig. 3).

**Figure 3.**
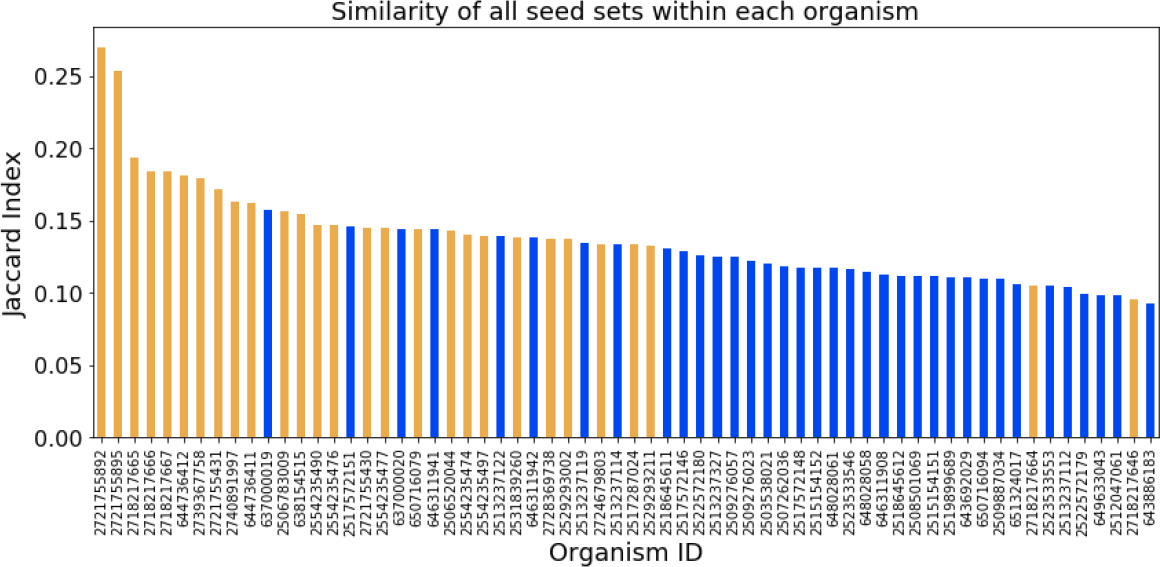
Similarity of all seed sets within each organism. The rank ordered mean jaccard index is shown for all 100 minimal seed sets we calculated for each organism. Bacteria are shown in blue and archaea are shown in orange.

We then turn to examine how seed sets necessary to produce viable organisms differ between organisms. We find that archaea seed sets tend to be more similar to one another than bacteria seed sets. Nonetheless, comparing organisms within domains leads to similar seed sets much more often than comparing organisms between domains (Fig. 4). This result holds true even when, instead of comparing the union of seed sets of organism 1 to the union of seed sets of organism 2, we compare the minimum seed set of organism 1 to organism 2. In this case we are looking at the minimal seed set of each organism that has the smallest mean molecular weight (Fig. 4B). However, we find that clustering the jaccard similarity between the union of organism seed sets results in more accurate clustering of the two domains we investigate (orange and blue squares above and to the left of the cluster maps show whether the row is an archaea or bacteria, respectively). The hierarchical clustering produced from unions shows that is is possible to correctly group archaea and bacteria from only their minimal seed sets necessary for viability. This is an interesting result, complementary to that of Ebenhoh et al (2006), who showed that organisms which are more closely related appear to have more similar reaction *scopes*, as measured by the Jaccard distance [6]. Such distinguishability in seed sets might be useful in identifying a relationship with taxonomy, for the purpose of expeditiously discerning the organisms which could be most likely to be risks for planetary contamination, or beneficial for terraformation.

**Figure 4.**
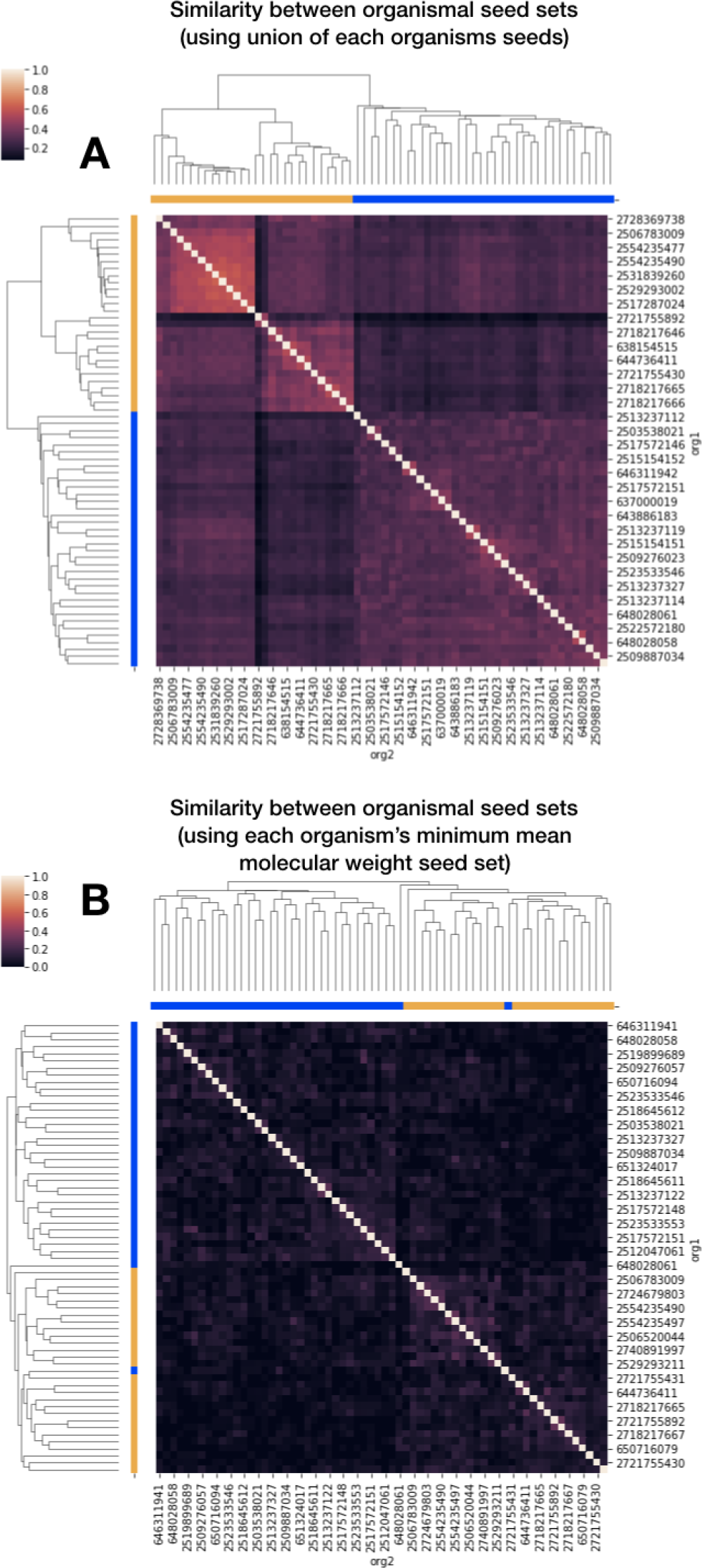
The similarity of seed sets between organisms. The clusters of two methods of organism comparisons are shown. (A) We take the union of all 100 seed sets within each organism, and compare them to one another using the jaccard index. (B) We take the minimal seed set of the smallest mean molecular weight of all 100 seed sets within each organism, and compare them to one another using the jaccard index. In both cases, the clustering separates out the domains (domain of each organism shown as blue squares for bacteria and orange squares for archaea above and below the cluster map.

**Figure 5.**
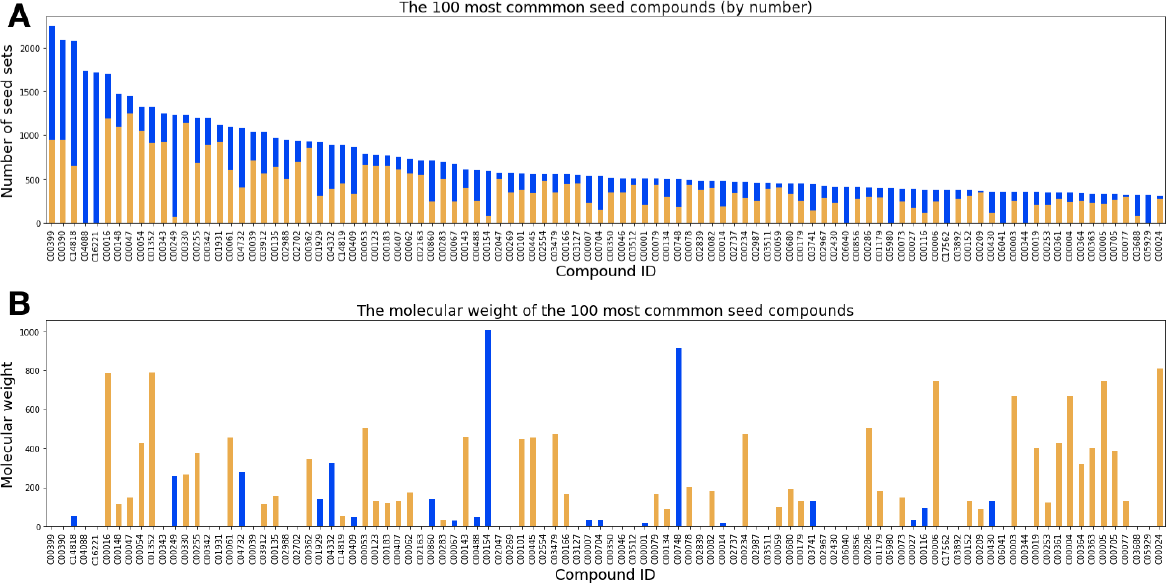
The top 100 most common seed compounds. (A) Rank ordered. The proportion of each found in archaea (orange) vs. bacteria (blue) seed sets are shown. (B) The molecular weights of each of the top 100 most common seed compounds. The domain of organism which most often contains seed sets with the compounds are shown as the color of the bar (archaea is orange and bacteria is blue).

**Figure 6.**
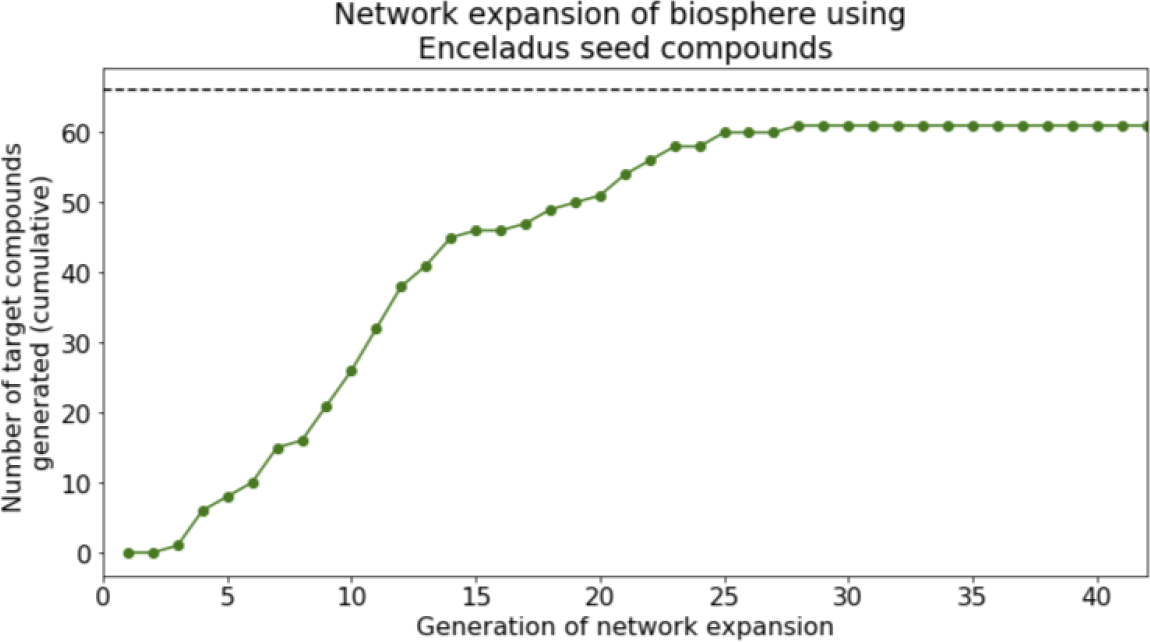
The network expansion of Earth’s biosphere using compounds available on Enceladus. Nearly all possible target metabolites are produced in this circumstance, with siroheme and heme being the notable missing compounds.

We turn to looking at the 100 most common seed compounds, to get some idea of the types of molecules we would expect to need to detect on Enceladus for this alkaliphiles to be viable. As might be expected, the majority of these compounds fall into common biochemical categories such as coenzymes, cofactors, amino acids, compound used for fatty acid synthesis, and other key metabolic pathways. It is notable that some of these compounds are target compound themselves, implying that these compounds are less likely to be synthesized by simpler compounds within these organismal metabolisms, and instead must be provided by the environment where possible.

Finally, we return to the initial set of seed compounds identified on Enceladus to examine if, with the full catalytic repoiroire of Earth’s biosphere utilizing the geochemistry of Enceladus, it is possible to produce the compounds essential for prokaryotic organismal viability. Using only the compounds identified on Enceladus, plus phosphate, leads to the ability to produce nearly all target metabolites, and those needed for most prokaryotic life. The expansion is missing siroheme, a cofactor used for sulfur reduction in metabolic pathways, as well as heme, a complex used for a variety of biological functions including electron transfer and redox reactions.

This would seem to indicate that if it was possible to transplant the entire catalytic repertoire of the Earth to Enceladus, it would be possible to maintain minimal metabolic viability for most prokaryotic organisms, provided that most of the reactions could be catalyzed in the high pressure alkaline environment. However, this is dependent on the exact structure of the individual organismal networks present. One strategy for terraforming might be to try and produce the minimal ecosystem which can reproduce the catalytic potential of the biosphere to send to another planet. Conversely, one potential strategy for making sure that a spacecraft is adequately sterilized might be to take a biological sample from a clean room spacecraft and annotate its metagenome. Then a network expansion could be run on the metagenomic network, with a conservative seed set, to ensure that none of the biochemistry would be viable at the spacecrafts destination.

## Discussion

In this research, we laid out a framework to quantify the chemical compounds necessary to assess the viability of Earth’s biochemistry in the context of other geospheres. We examine this framework as applied to Enceladus, executing the network expansion algorithm across metabolic networks of alkaliphilic bacteria and archaea in the chemical environment of Enceladus’ subsurface ocean. We find that no organisms analyzed can produce any of the pre-established target metabolites in this environment. However, a key element of life on Earth, phosphorous, has not yet been detected on Enceladus. We determine that incorporating phosphorous, by adding phosphate as a seed compound, does not change our results—there are still no target metabolites produced from any of the prokaryotes analyzed.

Next we investigated what it would take for these organisms to be viable, finding that the chemical complexity of the seed sets, or number of seeds present, has to be much higher. In many scenarios, both number of compounds and mean molecular weight of the compounds present must increase. By analyzing the jaccard index within minimal seed sets of organisms, we find that there are many unique seed sets which produce equally viable organisms across archaea and bacteria. We also find that between different taxa, seeds are more similar between two bacteria and between two archaea than when comparing organisms of different domains. The similarity of seed sets needed for organismal viability clusters organisms into their domains, indicating that there may be further ways to identify environments suitable to specific taxonomies across planets.

Finally we showed that when the catalytic capability of the entire biosphere is expanded around the Enceladus seed set (including phosphate), the target compounds necessary for viability are produced. This could indicate that, in principle, if the bulk biochemical diversity of Earth life could be transplanted to another planet via simple prokaryotic organisms, these organisms might be able to sustain a viable metabolism. Thus, embedding themselves into a planet from which they did not emerge with consequences for both life and the planet.

It is worth noting that the above study provides only a basic proof of concept for the idea of utilizing the well-developed technique of network expansion to quantitatively addressing the most pressing questions of astrobiology. There are many ways that this work could be expanded in order to better reflect geochemical reality as well as incorporate more theoretical considerations. For example, we could permute the initial conditions of the network expansions to force the inclusion of the observed Enceladus compounds into the randomized seed sets. Or we could more strictly constrain the shuffling between high/low molecular mass compounds in these seed sets.

There are further details which could help direct our search for compounds on other planets if we wish to improve this framework’s accuracy. For instance, we know that the presence/absence of cofactors is a big influence on the scope size for a seed set [13], so prioritizing our search for these compounds would provide high scientific returns. We could also measure viability as a gradient [8], and compare viability of organisms in other planetary contexts to the average viability of organisms across environments on Earth. We could investigate the specific metabolic pathways which are enriched or depleted in these environments [10]. Laboratory work here on Earth could also focus on better identifying reaction reversibility within organismal metabolic networks, as irreversible reaction networks would allow for more efficient algorithms used to identify minimal seed sets [2, 5, 15].

We might additionally included more statistical or theoretical constraints. For instance, can we identify distributions of molecular weights of compounds which tend to support biochemistry? Or link the expansions with knowledge of biochemical network topology, in order to find structural gaps in organismal networks which need to be filled to produce viable organisms [20]?

Moreover, the subset of organisms analyzed could be expanded to include organisms with greater metabolic diversity, or contracted to attempt to provide a better match between what we know about organismal environments on Earth with what we know about Enceladus. We could analyze the metabolisms of ecosystems, through metagenomic data, in addition to simple genomes. If we were specifically focused on the question of planetary contamination, we might also rerun these analyses on organisms which are known to exist in spacecraft sterilized clean rooms, like *Bacillus pumilus* SAFR-032 [36]. Further network expansion analyses could even be used to guide development of the composition of spacecraft materials to avoid metals which, if in contact with certain environments, could provide rich sources of cofactors or other compounds.

To summarize, the results from our network expansion analyses of alkaliphiles on Enceladus shows that there appears to be little risk of viability of these organisms, based on what we know about the chemical composition of the oceans. However, forward contamination, jeopardizing planetary protection, could be a much bigger risk if larger proportions of life’s catalytic potential are transported to other planets unintentionally. This seems remarkably less likely, although spacecraft clean room microbial ecosystems are not well characterized. Intentionally seeding a planet with life seems likely only in the circumstance where a metabolism is specifically tailored to the environment, and even then there are questions about how well it could be self-sustaining. We believe that because life on Earth was a product of Earth’s geochemistry, there is a significant bias to be viable only in a geochemical environment similar to the Earth’s. While there is much more work to be done to quantify the risks, or possibilities, of Earth life being viable amongst other geospheres, we believe that we have laid significant groundwork for exciting research in this domain.

## Materials and Methods

### Defining the networks

In order to run the network expansion algorithm from a seed set, we first had to define our networks. To identify the reactions and compounds present in the metabolic networks of individual organisms, we collected data from the Joint Genome Institute’s Integrated Microbial Genomes and Microbiomes database (JGI IMG/m) [26]. We located all archaea and bacteria which contained metadata on environmental pH, and filtered to those organisms with pH in the range of 9-11, approximately what might be expected in Enceladus’s ocean [9]. For our case study, we extracted data from all 28 archaea and 266 bacteria matching this criteria. We downloaded the Enzyme Commission (EC) numbers associated with each genome from the organism’s list of ‘Protein coding genes with enzymes‘. Each organisms list of EC numbers was mapped to the reactions which they catalyze using the Kyoto Encyclopedia of Genes and Genomes [16–18]. Using a combination of Biopython [4], the `KEGG REST API`, and `TogoWS` [19] to collect all KEGG `ENZYME, REACTION`, and `COMPOUND` data, we created reaction-compound networks for each organism. Each organisms network contains all of the reactions which all of its catalogued enzymes can catalyze, and all of the compounds involved in those reactions.

### Executing the network expansion

As outlined in the introduction, the network expansion process works as follows: An organism, defined by a fixed set of reactions which it has the ability to catalyze, can catalyze a reaction only if it has access to the necessary substrates. The initial substrates, called the seed set, are the compounds available to the organism from the environment. Initially, these are the only compounds in the organism’s network. The organism catalyzes all the reactions it can based on the reactions and compounds available in its network, and then adds the new compounds it can generate to its network. This process proceeds iteratively until the organism can produce no new compounds. The state of the organism’s network when expansion ceases is referred to as the organism’s scope—and it contains all of the compounds which can be synthesized by an organism, plus the seed set provided by the environment.

We assume that all reactions are reversible, both because the KEGG database recommends to not trust its reaction reversibility field, and because reaction directionality in nature depends on the concentrations of products and reactants, which we do not track here.

We ran the network expansion algorithm on the aforementioned subset of archaea and bacteria with documented environmental pH in the ranges of 9-11, using a seed set of compounds which have been identified on Enceladus from observations aboard CASSINI’s Ion and Neutral Mass Spectrometer (INMS) [37]. We additionally ran this seed set when including phosphate, which is likely present in small amounts from water-rock interactions, despite the lack of detection from Cassini’s INMS [11].

We also ran the network expansion of KEGG in its entirety (incorporating all catalogued compounds and reactions), representing the full catalytic and metabolic potential of the biosphere, on the seed set of Enceladus with phosphate (Table 2).

### Identifying minimal seed sets

We follow the algorithm described in Handorf et al., 2008 [12] to create random minimal seed sets which attempt to minimize the likelihood of obtaining seed sets with large complex biomolecules where possible:

A seed *S* is minimal if its scope Σ*S* contains the target compounds *T* and no proper subset of *S* fulfills this condition. *S* is a minimal seed set if:

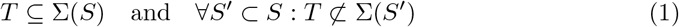

To find minimal seed sets for each organism, we start by creating a list of all the compounds involved in all the reactions that the organism can catalyze. Because the target compounds are by definition the intersection of an organisms compounds with the target metabolites, the target compounds must be present in this list. Going down the list, we check if removing a substrate will cause a network expansion seeded with the remaining substrates to successfully produce all target compounds. If the removal does not impact the target compounds produced, the substrate stays removed. Else, we add it back to the list. Then we move onto the next substrate in the list, repeating until the entire list is traversed.

In this algorithm, the order of the list affects the minimal seed set which gets identified, so it is necessary to permute the list and repeat the algorithm to identify each of the 100 minimal seeds. However, we do not want to start with a completely randomized list for each organism, because ideally we want to remove large complex compounds, as to be left with seed sets composed preferentially with simpler compounds which are more abiogenically plausible to find in a uninhabited environment. Previous research has shown that the scopes of single complex biochemicals tend to be reachable by sets of simpler molecules [13]. Because of this, we initially order every list from largest to smallest molecular weight, but then perturb them such that heavier compounds tend to stay near the top, thus getting preferentially removed. Compounds without associated weights were added in random locations in the list.

We again follow the method laid out by Handorf et al. [12]. From the list, two randomly chosen compounds with mass difference Δ*m* get exchanged with probability *p*:

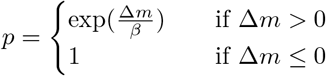

The only exception to this rule is that if one of the compounds does not contain weight information, then *p* = 0.5. The parameter beta represents the degree of disorder allowed in the list, where *β* = 0 forbids disorder and *β* = ∞ ignores disorder. We follow the choice of Handorf et al. [12] and choose *β* = 20 amu.

### Comparing and clustering seed sets

Similarity of seed sets were calculated using the Jaccard index. Clustering was computed using `scipy.cluster.hierarchy.linkage(method=’average’)`, where average refers to the unweighted pair group method with arithmetic mean (UPG-MA) algorithm.

## Acknowledgments

We thank the Emergence@ASU team (especially Doug Moore) for feedback and advice through various stages of this work.

